# The GHB analogue HOCPCA improves sensorimotor function after MCAO via CaMKIIα

**DOI:** 10.1101/2022.03.10.483849

**Authors:** Nane Griem-Krey, Anders B Klein, Bettina H Clausen, Mathias RJ Namini, Pernille V Nielsen, Heung-Chin Cheng, Cyrille Orset, Denis Vivien, Andrew N Clarkson, Kate L Lambertsen, Petrine Wellendorph

## Abstract

Ca^2+^/calmodulin-dependent protein kinase II alpha (CaMKIIα) is a major contributor to physiological and pathological glutamate-mediated Ca^2+^ signals, and its involvement in various critical cellular pathways demands specific pharmacological strategies. We recently presented GHB ligands as the first small molecules selectively targeting the CaMKIIα hub, a domain primarily responsible for holoenzyme oligomerisation, with an emerging functional role. Here, we report that the GHB ligand, HOCPCA, improves sensorimotor function after experimental stroke in mice when administered at clinically relevant time and in combination with alteplase. We observed that hub modulation by HOCPCA results in differential effects on distinct CaMKII pools, ultimately alleviating aberrant CaMKII signalling after cerebral ischemia. As such, HOCPCA normalised cytosolic Thr286 autophosphorylation after ischemia in mice and downregulated the ischemia-specific expression of a constitutively active CaMKII kinase fragment. Previous studies suggest holoenzyme stabilisation as a potential mechanism, yet a causal link to *in vivo* findings requires further studies. HOCPCA’s selectivity and absence of effects on physiological CaMKII signalling highlight pharmacological modulation of the CaMKIIα hub domain as an attractive neuroprotective strategy.

## Introduction

Even though ischemic stroke is one of the leading causes of death and disability,^1^ treatment options are limited. Thrombolysis using recombinant human tissue-type plasminogen activator (tPA, alteplase) or clot retrieval using mechanical thrombectomy have been shown to be effective in alleviating ischemic injury. However, only a small number of patients meets the criteria for treatment.^2,3^ Despite a great need for novel therapies targeting ischemic cell death mechanisms and enhancing functional outcome, so far, no cytoprotective therapy has successfully reached the market (reviewed in ^4^). Yet, positive results from a recent phase III trial (ESCAPE-NA1) bring new hope to the field, showing the feasibility of translating cytoprotective strategies to the clinic.^5^

The enzyme calcium/calmodulin-dependent kinase II alpha (CaMKIIα) is a major regulator of both physiological and pathological Ca^2+^ signals downstream of glutamate receptor activation, and has recently gained increasing attention as an attractive target for novel neuroprotective therapies.^6–10^ CaMKII has four subtypes, whereof CaMKIIα and CaMKIIβ are highly expressed in cerebral neurons forming large homo- and/or heteromeric structures made up of 12 to 14 subunits. Each subunit consists of a kinase, regulatory, linker, as well as a hub association domain.^11^ The enzyme is most known for its role in synaptic plasticity, where fine-tuned regulation by Ca^2+^ signals, multiple autophosphorylation sites as well as subcellular targeting enables a dual role of CaMKII in promoting both long-term potentiation and long-term depression (LTP and LTD).^12,13^ As such, elevated intracellular Ca^2+^ leads to stimulated activity mediated by Ca^2+^/CaM binding to the regulatory segment. Subsequent inter-subunit autophosphorylation at the central residue Thr286 (Thr287 in CaMKIIβ) infers Ca^2+^-independent autonomous activity, which is associated with translocation of the holoenzyme to the postsynaptic density (PSD) region in excitatory synapses, where it co-localizes with *N*-methyl-D-aspartate (NMDA) and α-amino-3-hydroxy-5-methyl-4-isoxazolepropionic acid (AMPA) receptors.^14,15^ Conversely, autophosphorylation of Thr305/306 (Thr306/307 in CaMKIIβ) can block Ca^2+^/CaM re-binding and inhibit further Ca^2+^-stimulated activity (reviewed in^16,17^).

During ischemia, intracellular Ca^2+^ overload leads to CaMKII dysregulation, as evidenced by an increase in autophosphorylation levels, long-lasting autonomous activity, and augmented PSD association.^8,12,19,20^ Furthermore, we recently reported the ischemia-specific cleavage of autophosphorylated CaMKII by calpain, releasing a constitutively active and stable CaMKII fragment (ΔCaMKII), which possibly contributes to enhanced substrate phosphorylation after ischemia.^20^ However, the individual contribution of these ischemic effects on CaMKII to cell death is unclear, and conflicting reports exist about CaMKII’s involvement in cell survival and cell death (reviewed in^21^). Remarkably, autonomous activity has been suggested as the main contributor to ischemic cell death based on strong neuroprotective effects with the peptide inhibitor, tatCN21.^6,7^ Yet, increased vulnerability to focal ischemia after conventional CaMKIIα knockout also indicates a role of CaMKII in cell survival.^8,22^ Due to CaMKII’s major role in cell signalling in both physiological and pathological conditions, there is a need to understand how CaMKII is involved in promoting cell death or survival to guide future pharmacological treatments.

We recently reported that pharmacological modulation of CaMKIIα by γ-hydroxybutyrate (GHB) and the GHB related small molecule, 3-hydroxycyclopent-1-enecarboxylic acid (HOCPCA), is neuroprotective when administered 30 min – 12 h after a photothrombotic stroke (PTS) in mice.^8,23^ GHB itself is a metabolite of the main inhibitory neurotransmitter γ-aminobutyric acid (GABA), and its neuroprotective properties have been described by several groups, however the mechanism has remained elusive.^24–26^ Whereas GHB also displays a low affinity for GABA^B^ receptors, possibly contributing to its neuroprotective actions via hypothermia^27^, its brain-penetrant analogue HOCPCA is endowed with remarkable selectivity and distinct affinity for the alpha subtype of CaMKII only, which is evident by a complete absence of binding of radiolabelled HOCPCA to brain tissue from *Camk2a*^*-/-*^ mice.^8,28,29^ In contrast to ‘classical’ CaMKII inhibitors,^30^ HOCPCA binds to a deep cavity of CaMKIIα’s hub domain.^8^ Its binding has no direct effect on substrate phosphorylation or Thr286 autophosphorylation, but instead leads to substantial stabilization of the oligomeric state of the hub domain when tested in a thermal shift assay.^8^ Changes in CaMKII oligomerisation upon HOCPCA binding are believed to alter holoenzyme functionality, yet the molecular mechanism and how this results in neuroprotection is unclear. Nonetheless, pharmacological modulation of the CaMKIIα hub domain presents an interesting therapeutic avenue, particularly because HOCPCA did not affect physiological CaMKIIα signalling, such as LTP.^8^

HOCPCA’s unique properties led us to characterize its neuroprotective potential further. Here, we present HOCPCA’s *in vivo* neuroprotective effect in two clinically relevant experimental stroke mouse models: the permanent middle cerebral artery occlusion (pMCAO) and thromboembolic stroke model. To investigate treatment effects on CaMKIIα biochemistry early after ischemia, we assessed CaMKII expression, autophosphorylation levels, as well as the presence of the cleavage product ΔCaMKII in membrane and cytosolic fractions after pMCAO.

## Results

### HOCPCA is neuroprotective after pMCAO

We have previously shown that a single dose of HOCPCA is neuroprotective when given 30 min and 3-12 h after PTS, both with effects on infarct size and motor function.^8^ As ischemic stroke is a clinically heterogeneous disease, it is recommended to study potential neuroprotectants in multiple models covering different disease aspects.^31,32^ Here, we tested HOCPCA after pMCAO, which shows a more pronounced ischemic penumbra compared to PTS.^33–35^ Hence, we initially treated mice intraperitoneal with 175 mg/kg HOCPCA or saline control early after pMCAO (30 min). Similar to Bach et al., neuroprotective outcomes were evaluated 3 days post-stroke by cresyl violet staining and grip strength test (summarized in **figure 1A**).^36^ HOCPCA treatment significantly reduced infarct volume by 26% compared to saline-treated mice (16.6 ± 5.9 mm^3^ for saline versus 12.3 ± 6.2 mm^3^ for HOCPCA, *p* = 0.0485; **Figure 1B**) and promoted functional recovery. Compared to baseline behavioural testing, pMCAO induced an asymmetry in the strength of the front paws (contralateral/ipsilateral; *p* = 0.0172; **Figure 1C**). This asymmetry was alleviated by HOCPCA treatment (**Figure 1C**). Intrigued by the neuroprotective effects of HOCPCA achieved with treatment 3-12 hours after PTS,^8^ we next tested if HOCPCA is also neuroprotective when given at a time when infarct development is known to be significantly progressed, i.e. 3 h post-pMCAO.^35^ Whereas such late treatment with HOCPCA did not influence infarct volume (19.5 ± 6.9 mm^3^ for saline versus 20.4 ± 7.8 mm^3^ for HOCPCA; **Figure 1D**), the effect on functional recovery persisted. As such, pMCAO introduced grip strength asymmetry in the control group (*p* = 0.0023), which was alleviated by HOCPCA treatment (**Figure 1E**).

**Figure 1:**
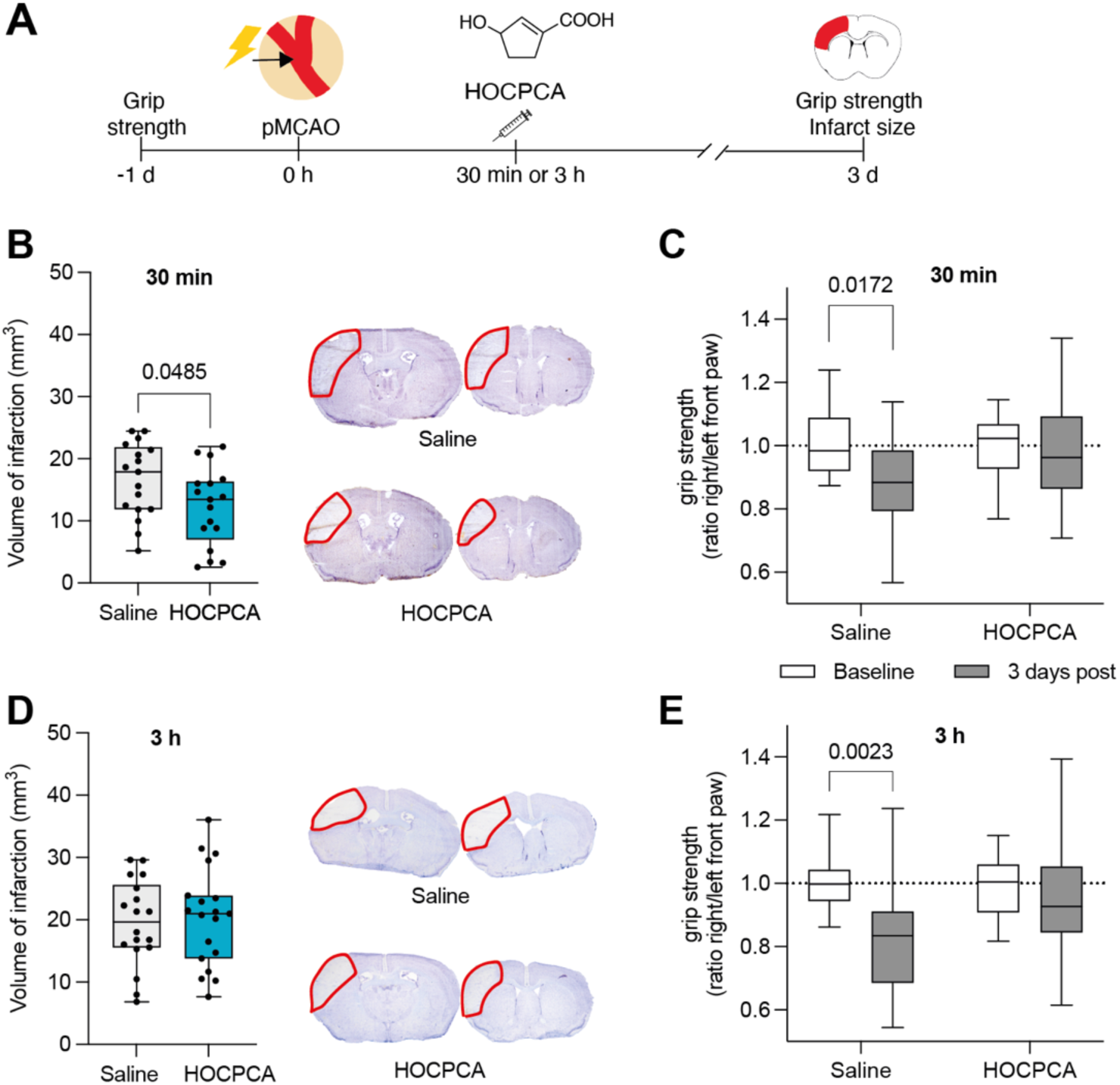
Effect of HOCPCA on infarct size and motor function after pMCAO. *(A)* Schematic illustration of experimental timeline. (*B-*E) Adult male mice were treated at 30 min (*B-C, n* = 17/group) or 3 h (*D-E, n* = 18-19/group) after pMCAO with 175 mg/kg HOCPCA or saline control (intraperitoneal). Infarct volumes were assessed 3 days post-stroke (*Left*), and representative cresyl violet stained tissue sections are shown (*Right*) (*B, D*; two-tailed Student’s *t* test). Forelimb motor function was evaluated 3 days post-stroke by grip strength analysis and shown as the ratio between right (contralateral) and left (ipsilateral) front paw (*C, E*; two-way ANOVA (time, treatment), post hoc Tukey’s test) (Box plot (boxes, 25–75%; whiskers, minimum and maximum; lines, median).

### HOCPCA reverses pThr286 levels in the cytosolic fraction of peri-infarct cortex after stroke

Prompted by HOCPCA’s neuroprotective effects after pMCAO, we next investigated the potential mechanism of action *in vivo*. Given HOCPCA’s well-characterized binding to the CaMKIIα hub domain,^8^ an allosteric effect was envisaged. However, little is known about HOCPCA’s biochemical effects on CaMKIIα signalling. After an ischemic insult, CaMKIIα is rapidly activated, which is resulting in increased autophosphorylation and translocation to the PSD.^37^ We thus initially evaluated HOCPCA’s effect on Thr286 autophosphorylation after stroke. As we saw a robust neuroprotective effect of HOCPCA in this model 30 min post-pMCAO (**Figure 1A**), we chose this time point for treatment and harvested peri-infarct tissue 2 h post-stroke (**Figure 2A**), which is reported as a suitable time point to detect CaMKII biochemical alterations after MCAO.^19,37^ Surprisingly, we did not observe any effect of stroke or HOCPCA treatment on Thr286 autophosphorylation in homogenates from the peri-infarct cortex (**Suppl. Figure 1**).

**Figure 2:**
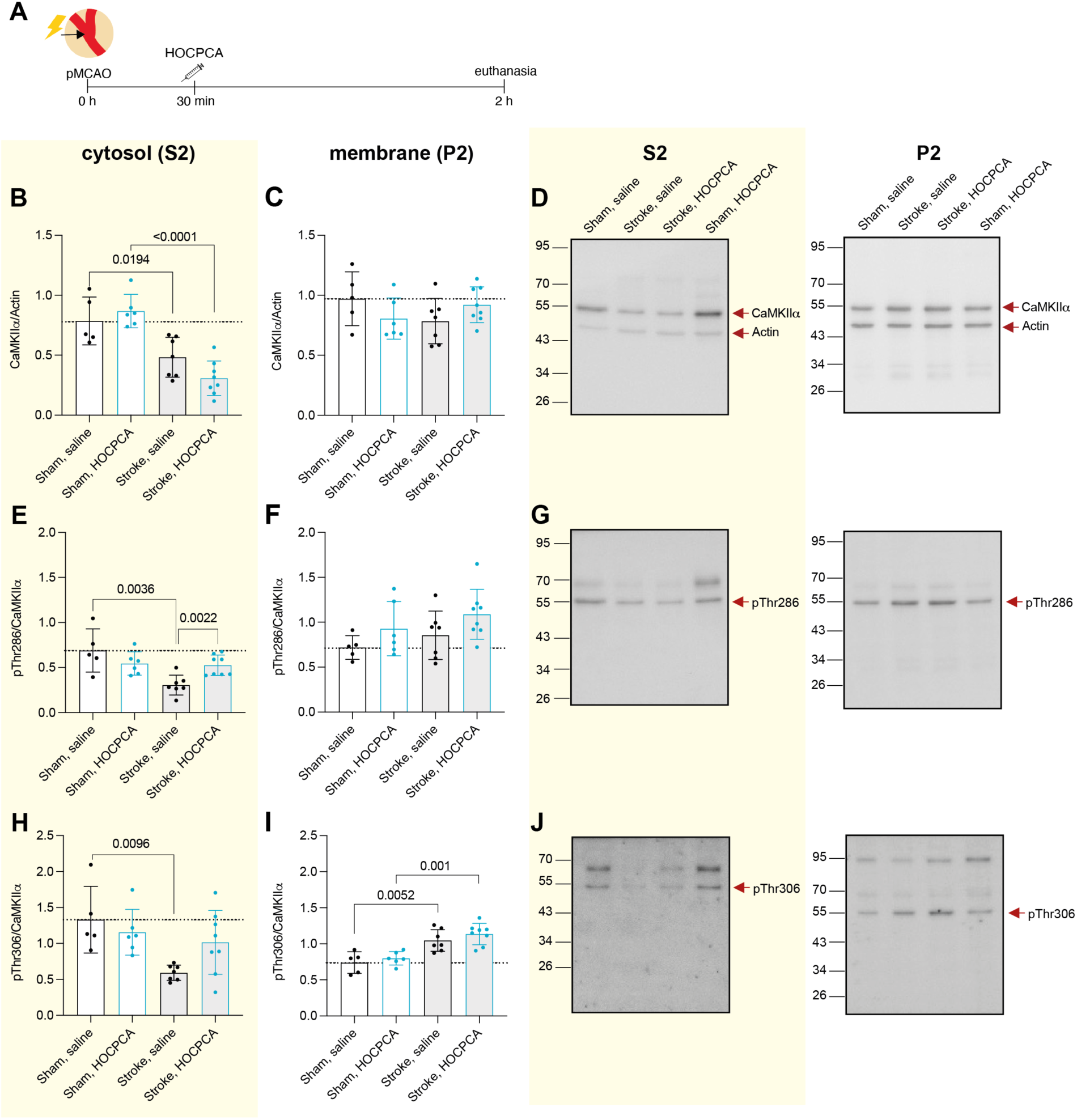
Effect of HOCPCA on CaMKIIα expression and autophosphorylation in subcellular fractions after pMCAO. *(A)* Schematic illustration of experimental timeline. Adult male mice were subjected to sham (*n* = 5-6/group) or pMCAO-surgery (*n* = 7-8/group), treated with 175 mg/kg HOCPCA or saline control, and brains were harvested 2 h post-stroke. (*B-*J) Membrane (P2) and cytosolic (S2) fractions of homogenates from peri-infarct tissue were probed for total CaMKIIα (*B-D*), pThr286 (*E-G*), or pThr306 (*H-J*) and normalized to Actin or total CaMKIIα expression. (*D, G, J*) Representative Western blots. (*Note:* Western blots were performed blinded, and the order of treatment groups differs in bar diagrams for most logical interpretation; Full blots are shown, and three technical repetitions were performed; One-way ANOVA, post hoc Tukey’s test, only significance for the following comparisons is presented in the graphs for simplification: sham, saline versus stroke, saline; sham, HOCPCA versus stroke, HOCPCA, and stroke, saline versus stroke, HOCPCA).

After ischemia-induced translocation to the PSD, CaMKIIα interacts with the NMDA receptor subunit GluN2B.^38^ Because HOCPCA reduces this colocalization *in vitro*,^8^ we investigated the effect on total CaMKIIα expression in both membrane (P2) and cytosolic (S2) subcellular fractions after pMCAO. Consistent with previous reports,^19^ we observed that total CaMKIIα expression significantly decreased in the cytosolic S2 fraction after pMCAO compared to sham (*p* = 0.0194). However, we did not observe any effect of HOCPCA on total CaMKIIα expression in sham or stroke injured tissue (**Figure 2B**). As CaMKIIα is differently autophosphorylated depending on its subcellular location,^6^ we also evaluated its autophosphorylation levels in these subcellular fractions. We observed a decrease in pThr286 after pMCAO in the cytosolic S2 fraction (*p* = 0.0036). Interestingly, HOCPCA treatment significantly restored Thr286 phosphorylation to sham levels (66% increase, *p* = 0.002; **Figure 2E**). A similar, albeit smaller non-significant decrease was seen for pThr306 autophosphorylation (*p* = 0.1329; **Figure 2H**).

No stroke or treatment effect was observed on total CaMKIIα as well as pThr286 expression in the membrane P2 fraction (**Figure 2C, F**). However, we observed increased pThr306 levels after pMCAO in the membrane P2 fraction, both in saline- and HOCPCA-treated mice (**Figure 2I**).

### HOCPCA does not affect pThr287-CaMKIIβ in the cytosolic fraction after pMCAO

As CaMKIIα assembles with CaMKIIβ into mixed holoenzymes with a 3:1 ratio within forebrain regions,^39^ we also evaluated whether HOCPCA binding to CaMKIIα could affect CaMKIIβ activity after pMCAO. Similar to CaMKIIα, total CaMKIIβ expression was significantly decreased in the cytosolic fraction after pMCAO, and no effect of HOCPCA treatment was observed (**Figure 3A**). However, in contrast to CaMKIIα, corresponding Thr287 autophosphorylation of CaMKIIβ was neither affected by pMCAO or HOCPCA treatment (**Figure 3D**). This underlines the alpha subtype selectivity of HOCPCA, also *in vivo*. Moreover, whereas we did not observe effects of ischemia or treatment on total and pThr286 expression of CaMKIIα in the membrane P2 fraction, pMCAO resulted in an increase in total CaMKIIβ and decrease in pThr287 expression within the P2 fraction (**Figure 3B, E**) in the HOCPCA treated samples only.

**Figure 3:**
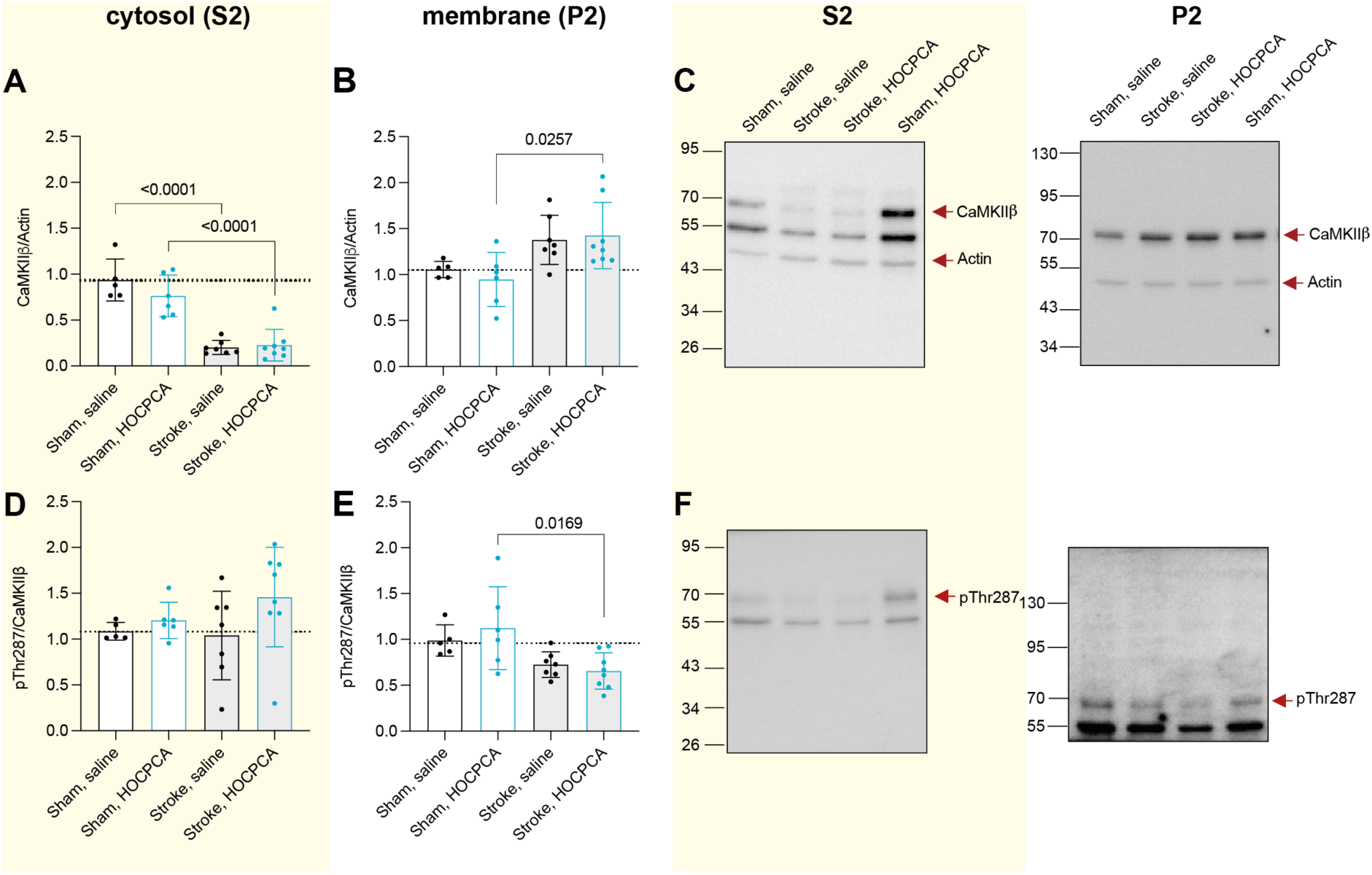
Effect of HOCPCA on CaMKIIβ expression and autophosphorylation in subcellular fractions after pMCAO. (*A-*F) Membrane (P2) and cytosolic (S2) fractions of homogenates from peri-infarct tissue from sham (*n* = 5-6/group) or pMCAO-operated mice (*n* = 7-8/group) were probed for total CaMKIIβ (*A-C*) and pThr287 (*D-F*). Expression was normalized to Actin or total CaMKIIβ expression, respectively. (*C,F*) Representative Western blots. (*Note:* Western blots were performed blinded, and the order of treatment groups differs in bar diagrams for most logical interpretation; Full blots are shown, and three technical repetitions were performed; One-way ANOVA, post hoc Tukey’s test, only significance for the following comparisons is presented in the graphs for simplification: sham, saline versus stroke, saline; sham, HOCPCA versus stroke, HOCPCA, and stroke, saline versus stroke, HOCPCA). Experimental details are the same as in Figure 2.

### HOCPCA downregulates an ischemia-specific cleavage fragment of CaMKII in the membrane fraction

We have previously reported the ischemia-induced cleavage of both Thr286 autophosphorylated CaMKIIα as well as Thr287 autophosphorylated CaMKIIβ by calpain after PTS.^20^ Specifically, we observed a 31 kDa-fragment of CaMKII (referred to as ΔCaMKII) after stroke, which corresponds to the kinase domain devoid of its regulatory, linker, and hub domain, rendering the kinase constitutively active. We suggest that this contributes to ischemia-induced CaMKII dysregulation (**Figure 4A**),^20^ and that HOCPCA influences CaMKII cleavage after ischemia via a unique molecular interaction with the hub. To investigate this, we initially investigated whether ΔCaMKII was present in the subcellular pMCAO samples. To this end, we probed with a CaMKIIpan antibody, which detects an epitope within the highly conserved kinase domain. Hence, the antibody does not discriminate between CaMKII subtypes, and the ΔCaMKII might be derived from either CaMKIIα or CaMKIIβ. As in the PTS model,^20^ we saw a dramatic increase in ΔCaMKII expression in the membrane P2 fraction after pMCAO (230% increase compared to sham; *p* = 0.006; **Figure 4B**). Remarkably, in the HOCPCA-treated samples this was significantly reduced (48% decrease compared to pMCAO; *p* = 0.0067), thus indicating a treatment effect on the ischemia-specific expression of βCaMKII. We were unable to detect ΔCaMKII in the cytosolic S2 fraction.

**Figure 4:**
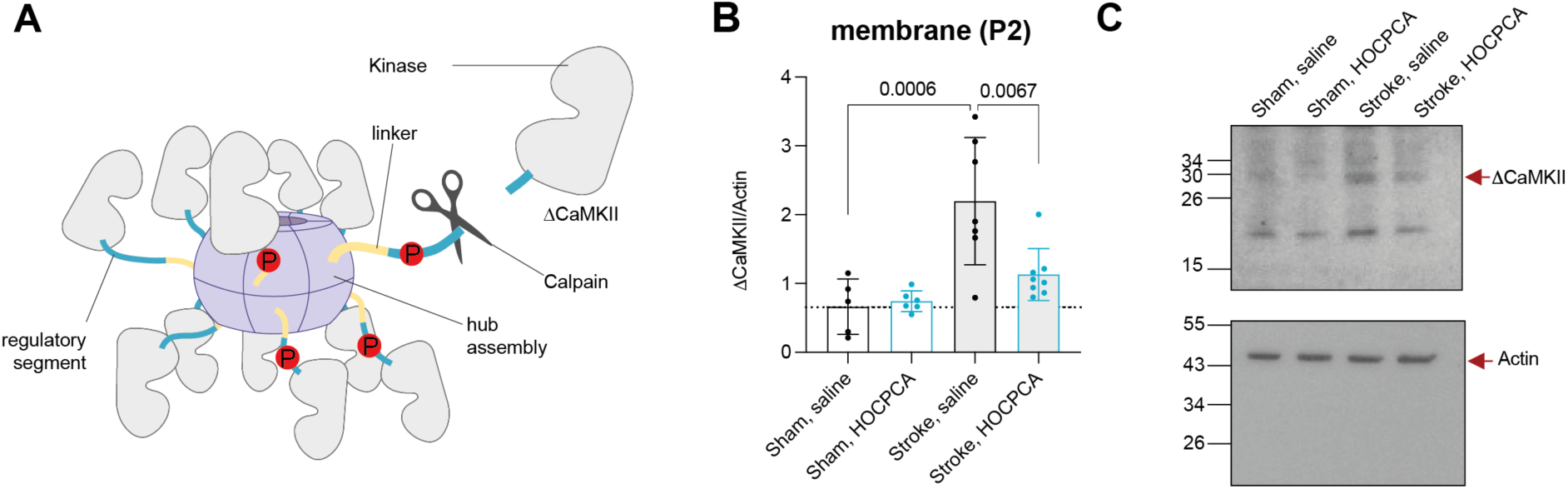
Effect of HOCPCA on CaMKII cleavage after pMCAO. *(A)* Graphical illustration of CaMKII cleavage after ischemia. *(B)* A 31 kDa N-terminal fragment of CaMKII (βCaMKII) was detected by CaMKIIpan antibody in the P2 membrane fraction after pMCAO (*n* = 7-8/group) or sham surgery (*n* = 5-6/group), and expression was normalized to Actin (see figure 2C and 3B for total CaMKIIα and CaMKIIβ expression) (*C*) Representative Western blots. (Western blots were performed blinded; Full blots are shown, and three technical repetitions were performed; One-way ANOVA, post hoc Tukey’s test, only significance for the following comparisons is presented in the graphs for simplification: sham, saline versus stroke, saline; sham, HOCPCA versus stroke, HOCPCA, and stroke, saline versus stroke, HOCPCA). Experimental details are the same as in Figure 1.

### HOCPCA improves motor function after thromboembolic stroke

To extend the clinical translatability, we further assessed HOCPCA’s neuroprotective effect in a mouse model with a reperfusion component, in this case a thromboembolic stroke. Here, *in situ* injection of thrombin leads to MCAO by the local formation of a fibrin-rich clot, thus enabling the study of neuroprotective agents in combination with tPA-induced reperfusion.^40^ The model is uniquely characterized by gradual reperfusion hours after thrombin injection mediated by spontaneous clot dissolution by the endogenous fibrinolytic system,^41,42^ and thus resembles human pathology more closely compared to classical mechanical transient MCAO where blood flow is reversed promptly and with a much larger lesion.^35^

To our knowledge, ischemia-induced CaMKIIα dysregulation has never been addressed in this model. Therefore, we initially examined pThr286 levels as a measure of CaMKIIα dysregulation. A time point 30 min post-stroke was selected as this represents the desired time for HOCPCA treatment (**Figure 5A**). Compared to sham, we found that pThr286 levels were significantly increased after stroke while total CaMKIIα expression remained unchanged (**Figure 5B**), confirming pathological CaMKIIα activation also in this model. Subsequently, we went on to test HOCPCA’s neuroprotective effect on infarct size and motor function. To this end, mice received either a single dose of 175 mg/kg HOCPCA at 30 min, intravenous administration of 10 mg/kg tPA starting 20 min post-thrombin injection, saline control, or a combination of HOCPCA and tPA treatment (**Figure 5A**). Infarct volumes were evaluated 1-day post-stroke and evaluated by T2-weighted imaging. While tPA treatment significantly reduced infarct size by 39% (23.7±7.2mm^3^ for saline versus 14.4±5.8mm^3^ for tPA; *p* = 0.0035), which is similar to previous reports,^43^ neither HOCPCA treatment alone nor a combination of tPA and HOCPCA treatment affected infarct size (23.3±7.7mm^3^ for HOCPCA and 17.6±7.2mm^3^ for HOCPCA+ tPA; **Figure 5C**). By contrast, we could detect neuroprotective effects of all treatment regimens on functional recovery. Specifically, HOCPCA, tPA as well as HOCPCA+ tPA treatment relieved stroke-induced forelimb asymmetry in grip strength when assessed 3 days post-stroke (**Figure 5D**). Only saline-treated mice showed forelimb grip strength asymmetry (*p* = 0.0009 at day 1 and *p* = 0.0254 at day 3), similar to previous reports in this model.^44^ Moreover, the protective effect of HOCPCA treatment on grip strength was already seen at day 1 post-stroke, whereas this was not the case for tPA (*p* = 0.0145) or HOCPCA+ tPA (*p* = 0.0064) groups (**Figure 5D**). Finally, to further characterize HOCPCA’s neuroprotective effect, we evaluated if early HOCPCA treatment affects the gene expression of selected inflammatory markers. To this end, we evaluated mRNA expression of the cytokine TNFα and the microglia/macrophage activation markers Iba1 and CD68 in the ischemic cortex 3 days after both thromboembolic stroke and pMCAO. HOCPCA treatment was able to reduce mRNA expression of inflammatory markers compared to saline in both models. Similar effects were seen for tPA and tPA + HOCPCA treatment after thromboembolic stroke (**Suppl. Figure 2**).

**Figure 5:**
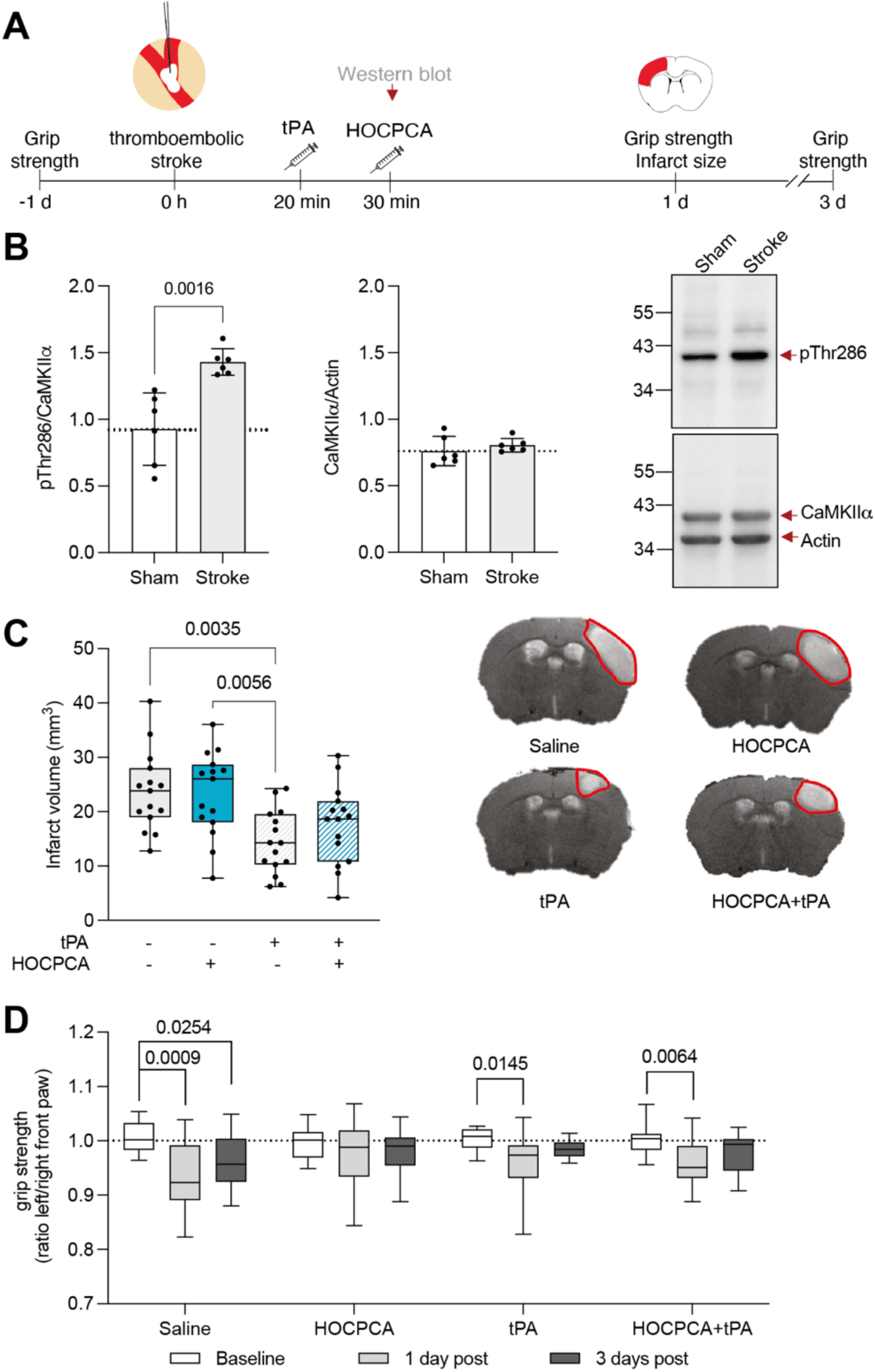
Validation of HOCPCA’s neuroprotective effect using a thromboembolic stroke model with reperfusion component. *(A)* Schematic illustration of experimental timeline. *(B)* CaMKIIα pThr286 autophosphorylation and total CaMKIIα expression 30 min post-thromboembolic stroke compared to sham (*n* = 6/group). Expression was normalized to total CaMKIIα or Actin, respectively. Representative Western blots are shown to the right. (*C,D*) Adult male mice were treated either with saline, tPA (intravenous, 10 mg/kg, 10% bolus/90% infusion over 40 minutes starting from 20 min after occlusion), HOCPCA (intraperitoneal, at 30 min post-stroke) or a combination of tPA and HOCPCA (*n* = 15/group). (*C*) Infarct size was assessed by by T2-weighted imaging 1-day post-stroke (One-way ANONA, post hoc Tukey’s test). (*Right*) Representative by T2-weighted images. (*D*) Grip strength represented as the ratio between left (contralateral) and right (ipsilateral) front paw was assessed 1 day and 3 days after stroke (two-way ANOVA (time, treatment), post hoc Tukey’s test) (Box plot (boxes, 25–75%; whiskers, minimum and maximum; lines, median).

## Discussion

CaMKIIα is a major mediator of physiological and pathological Ca^2+^ signals regulating numerous downstream pathways with possible roles in cell death or cell survival (reviewed in ^21^). Moreover, ischemia leads to CaMKII dysregulation in various ways. For instance, subsets of CaMKII with increased autonomous activity, elevated levels of multiple autophosphorylation sites, PSD translocation as well as calpain-mediated cleavage into a stable constitutively active kinase fragment have been described.^6, 19,45^ Yet, the exact interplay and which mechanism constitutes the best pharmaceutical target is still to be fully elucidated and warrants much needed further studies.

### HOCPCA treatment is neuroprotective and reverses ischemia-induced CaMKII dysregulation

In this study, we establish pharmacological modulation of the CaMKIIα hub domain by GHB ligands as an attractive strategy to reduce CaMKII dysregulation after stroke in mice. First, we show that the GHB analogue, HOCPCA, improves motor function in two clinically relevant murine stroke models and is able to dampen inflammatory changes. Second, we present unforeseen and differential effects of HOCPCA treatment on distinct subsets of ischemia-dysregulated CaMKII pools. Yet, elucidating the underlying molecular mechanisms mediated by ligand binding to the hub domain requires further studies.

HOCPCA reversed an ischemia-induced decrease in cytosolic Thr286 autophosphorylation. Ischemia-induced translocation of the cytosolic CaMKIIα has been previously reported to coincide with a further decrease in cytosolic pThr286.^6^ Yet previous studies focused on the effects of CaMKII dysregulation at the PSD, where increased pThr286 as well as autonomous activity were shown to be associated with death promoting CaMKII effects.^6^ Collectively, this might indicate that the previously described dual role of CaMKIIα during ischemia is differentiated by pThr286 levels in distinct subcellular locations, i.e., membrane-anchored pThr286 CaMKIIα might promote cell death, whereas normalization of pThr286 levels within the cytosolic compartment might instead lead to neuroprotection.

Moreover, we show that HOCPCA reduces ΔCaMKII expression levels in the membrane fraction, which constitutes an ischemia-specific stable fragment of CaMKII.^20^ Strikingly, we recently documented that calpain mediates interdomain cleavage of Thr286 autophosphorylated CaMKIIα right before the regulatory segment, thus releasing a kinase domain fragment (ΔCaMKII) that is by nature constitutively active.^20^ In fact, ΔCaMKII has been widely used as a tool to study the impact of CaMKII activity,^46–49^ yet, its natural occurrence after pMCAO has never been reported and highlights it as a potential pathospecific target in excitotoxicity. Interestingly, we detected ΔCaMKII only in the membrane fraction and not in the cytosol, supporting a membrane-associated death-promoting role for CaMKIIα. Of note, the binding site for GluN2B subunit of the NMDA receptor is mapped to the kinase domain of CaMKIIα,^14^ thus suggesting that membrane anchoring of ΔCaMKII is still feasible. Mechanistically, calpain-mediated cleavage of pThr286-CaMKIIα might act as a downstream neurotoxic effect of the sustained ischemia-induced increase in pThr286.^6,37^ As such, unregulated kinase activity of βCaMKII is likely contributing to excessive substrate phosphorylation after ischemia, resulting in ischemic cell death. Most interestingly, HOCPCA treatment reduced the expression of βCaMKII after stroke, suggesting an ischemia-specific compound effect that might be involved in the reversal of CaMKII dysregulation. Of note, even though calpain inhibition has been shown to be neuroprotective,^50^ the exact role of βCaMKII in neuronal cell death as well as the mechanism of HOCPCA-mediated reduction in ΔCaMKII remains to be determined. Still, it offers an attractive new way of salvaging tissue damage after stroke.

### HOCPCA acts selectively and specifically under ischemic conditions

HOCPCA is part of a new class of CaMKIIα ligands (GHB analogues) with unparalleled selectivity for the hub domain of the alpha subtype. While ‘classical’ CaMKII inhibitors such as, KN93 and CN-peptides, not only show off-target effects, they also inhibit all four CaMKII subtypes.^51,52^ This increases the risk for unwanted side effects as CaMKII subtypes are expressed throughout all tissues. In fact, CaMKIIδ plays a major role in cardiomyocyte signalling, however, emerging evidence is also showing a role for CaMKIIδ in regulating astrocyte and neuronal cross-communication and cell survival after ischemia.^53–55^ Moreover, the use of CaMKII inhibitors might be hampered due to possible effects on learning and memory. As such, tatCN21 interfered with learning ability in mice by inhibiting LTP induction.^56^ In contrast, HOCPCA did not affect LTP, and most intriguingly, we did not observe effects of HOCPCA on CaMKIIα during naïve nonpathological conditions.^8^ Accordingly, HOCPCA was only neuroprotective when applied after a glutamate insult in primary cortical neurons, thus indicating the need for CaMKII dysregulation for effective hub modulation.^8^ In addition, HOCPCA selectively normalized cytosolic pTh286-CaMKIIα after stroke, while no treatment effect was seen in the membrane fraction nor in naïve mice or on pTh287-CaMKIIβ, supporting its selectivity and pathospecific action *in vivo*. The observed effect on pThr286 is in contrast to our previous finding that HOCPCA did not affect pThr286 in basal or Ca^2+^-stimulated cortical primary neurons.^8^ Yet, a potential effect of HOCPCA on pThr286 after Ca^2+^-stimulation was likely masked by studying whole cell lysates instead of subcellular fractions, similarly we did not observe any effect of HOCPCA on pThr286 in whole-cell homogenates in this study.

### Mechanistic thoughts behind HOCPCAs effects in peri-infarct subcellular compartments

At this point, it is unclear how pharmacological modulation of the hub domain results in the normalization of cytosolic pThr286 levels, yet an allosteric effect via the hub domain is envisaged. In fact, recent reports highlight an emerging functional importance of the hub domain. For instance, a role in allosteric modulation of kinase activity^57^ as well as in activation-triggered destabilization of holoenzymes and subsequent dimer release and subunit exchange have been suggested.^58–60^ Interestingly, we previously showed that HOCPCA is able to stabilize CaMKIIα hub oligomerization *in vitro* in a thermal shift assay.^8^ Although a causal mechanistic link is absent, it is tempting to speculate that the ischemia-induced restorative effect of HOCPCA on pThr286 levels within the cytosolic fraction could be mediated by increased holoenzyme stability possibly associated with a reduction in activation-triggered subunit release. Specifically, as pThr286 occurs in trans between neighbouring activated subunits,^61^ stabilization of the oligomeric state of the holoenzyme could be apparent in increased Thr286 autophosphorylation. Yet, other mechanisms could also be involved, such as differential regulated phosphatase activity.^62^

Moreover, we did not observe increased pThr286 or total CaMKIIα expression, nor a treatment effect of HOCPCA in the P2 membrane fraction 2 h after pMCAO in this study. This is in contrast to our previous finding that HOCPCA inhibited glutamate-induced CaMKIIa-GluN2B colocalization in primary hippocampal neurons^8^ and to several groups reporting sustained ischemia-induced CaMKII translocation and autophosphorylation at the PSD.^6, 19,37^ Yet, the P2 fractions generated in this study include the enrichment of total cell membranes. PSD or synaptic/extrasynaptic fractions might have been better suited for the accurate detection of CaMKIIα translocation. Nonetheless, the discrepancy could also be explained by inherent differences in tissue and/or injury-specific CaMKIIa responses. As such, dysregulated CaMKII translocation and autophosphorylation are well characterized in hippocampal tissue after global ischemia^6, 18,63–66^ and cortical tissue after transient MCAO.^19, 37,67^ CaMKIIα’s subcellular location after pMCAO might involve differential kinetics compared to previous studies, including injury with a reperfusion component. This is supported by data from Skelding et al., who studied the timing of pThr286 after transient MCAO before and after reperfusion.^68^ Skelding and colleagues observed an initial increase in pThr286, which was downregulated at 90 min after MCAO, and then subsequently after reperfusion, a second Ca^2+^ wave triggered another increase in pTh286, indicating a pattern of phosphorylation and dephosphorylation depending on intracellular Ca^2+^ levels. ^68^ Intriguingly, the same study did not detect an effect of ischemia on pThr305/306 in homogenates, whereas our results indicated an ischemia-induced increase in pThr305/306 in the P2 membrane fraction. As pThr305/306 is located within the CaM binding element, this could indicate dissociation of Ca^2+^/CaM from CaMKIIα, which may further support the hypothesis that 2 h after pMCAO the initial Ca^2+^ wave is ceased. Nonetheless, CaMKIIα dysregulation might still be sustained at this time point, and yet manifested in a way other than pThr286 activation. This is supported by the fact that HOCPCA treatment 3 h post-pMCAO could still improve functional outcome.

### HOCPCA is neuroprotective in multiple in vivo stroke models

Until this point, we have observed neuroprotective effects of HOCPCA in three different animal models of stroke performed in separate laboratories (this study and Leurs et al.).^8^ Not surprisingly, the therapeutic window and effect size was considerably different between models, which might be explained by their different aetiology.^34,69^ Of note, HOCPCA was able to improve motor function after thromboembolic stroke in combination with tPA, suggesting no negative interaction of co-administration as observed in the ESCAPE-NA1 trial.^4,5^

Nonetheless, HOCPCA treatment 30 min post-stroke was less effective after thromboembolic stroke compared to pMCAO. Interestingly, similar to the human pathology, this model is characterized by spontaneous reperfusion.^41,42,70^ As shown by Skelding et al.,^68^ reperfusion could lead to a second Ca^2+^ wave and CaMKIIα dysregulation, possibly further aggravating the injury. As HOCPCA is endowed with a short half-life of 20 min,^28^ its effect might have been dampened by secondary injury or CaMKII activation caused by reperfusion. Consequently, it would be interesting to see if a second dose after reperfusion could lead to effects on infarct size in that model. Nonetheless, the reduced effect of HOCPCA in the thromboembolic model compared to pMCAO could also be caused by differences in anaesthesia type and length (inhalation versus general anaesthesia). As such, the anaesthetics might differently regulate blood pressure, cerebral blood flow or have neuroprotective properties itself, potentially affecting the neuroprotective outcome.^34^ Finally, HOCPCA’s ability to dampen inflammatory changes substantiates its effects on functional outcome after stroke, yet, the underlying mechanisms require further investigation.

## Conclusion

In this study, we present pharmacological modulation of the CaMKIIα hub domain by HOCPCA as a strong therapeutic candidate for ischemic stroke by improving functional outcome in multiple stroke models via a novel mechanism-of-action. As such, HOCPCA treatment had differential effects on distinct subcellular pools of CaMKIIα, thereby alleviating ischemia-induced CaMKII dysregulation, i.e., HOCPCA normalized pThr286 in the cytosol and downregulated ΔCaMKII in the membrane fraction. Even though previous studies suggest holoenzyme stabilization as a potential underlying mechanism, further studies are needed to establish a causal link and to decipher downstream functional consequences. Yet, HOCPCA’s high degree of selectivity, pathospecific mechanism as well as the neuroprotective effect in multiple *in vivo* mouse models shows that the brain-penetrable small-molecule shows great potential. Further understanding of both temporal and mechanistic CaMKII dysregulation after different ischemic insults will direct future pharmacological treatment regimes.

## Acknowledgments

This work was supported by the Lundbeck Foundation (R277-2018-260) and the Novo Nordisk Foundation (NNF14CC0001). The Emil Herborg Grant and the Julie von Müllen Foundation supported the work with travel grants. We thank Bente Frølund for providing HOCPCA. A special thanks to Florent Lebrun for his help with the thromboembolic stroke surgery. We thank Diana Wichmann Graff, Monica Santiveri-Saez and Carina Bachmann for their help with stroke experiments.

## Author contribution statement

NGK, ABK, BHC, CO, DV, ANC, KLL and PW designed the studies. NGK, ABK, BHC, PVN and KLL performed and analysed pMCAO *in vivo* studies, whereof BHC performed grip strength test. NGK and OC performed and analysed thromboembolic stroke and grip strength studies. NGK and MRJ performed and analysed subcellular fractions, Western blot and qPCR studies. HCC conceptualized and assisted with CaMKII cleavage studies. The manuscript was written by NGK and revised by PW. All authors, reviewed, edited, and approved the final version of the manuscript.

## Materials and methods

### 1. Materials

The sodium salt of HOCPCA (3-hydroxycyclopent-1-enecarboxylic acid) was synthesized as previously described.^29^ Mouse α-thrombin (#90792 MIIA) was obtained from Stago BNL (the Netherlands) and tissue-type plasminogen activator (tPA, Actilyse®) from Boehringer Ingelheim (Germany).

The following antibodies have been validated in Leurs et al. with *Camk2a -/-* and *Camk2b -/-* tissue: CaMKIIα (#NB100-1983, RRID:AB_10001339; mouse monoclonal IgG, clone 6g9, Novus Biologicals; 1:10 000 overnight incubation at 4 °C, stored at -20 °C), CaMKIIβ (#12716, RRID:AB_2713889; mouse monoclonal IgG2b, clone CB-beta-1, Invitrogen; 1:5000 overnight incubation at 4 °C, stored at -20 °C), phospho-CaMKII (Thr286/287) (#AP12716b, RRID:AB_10820669; rabbit monoclonal IgG, clone D21E4, Cell Signaling Technology; 1:1000 overnight incubation at 4 °C, stored at -20 °C).^8^ In addition, the following antibodies have been used: CaMKII (pan) (#4436, RRID:AB_1054545; rabbit monoclonal IgG, clone D11A10, Cell Signaling Technology; validated in Ameen et al.;^20^ 1:500 overnight incubation at 4 °C, stored at - 20 °C), Phospho-CaMKII (Thr306) (#p1005-306, RRID: AB_2560940, rabbit polyclonal IgG, 1:1000 overnight incubation at 4 °C, stored at -20 °C), beta-Actin (#ab6276, RRID: AB_2223210; mouse monoclonal IgG, cloneAC-15, Abcam, validated by company (https://www.abcam.com/beta-actin-antibody-ac-15-ab6276.html#lb); 1:10 000 1 h incubation at room temperature, stored at -20 °C). The following secondary antibodies have been used: Goat anti-mouse Alexa Fluor Plus 800 (#A32730, RRID: AB_2633279; polyclonal IgG, Invitrogen; 1:2000 1 h incubation at room temperature, stored at 4 °C) and donkey anti-rabbit Alexa Fluor Plus 488 (#A32790, RRID: AB_2762833; polyclonal IgG, Invitrogen; 1:5000 1 h incubation at room temperature, stored at 4 °C).

### 2. Animals

All animal experiments were approved by the individual ethics committees according to national and international guidelines and are reported after the ARRIVE guidelines (Animal Research: Reporting In Vivo Experiments). Mice were allowed to acclimatize for at least one week before experiments and were allocated to treatment groups by random. The experimenter was blinded to treatment groups during allocation, conduction of experiments as well as data analysis. Mice were excluded if the stroke surgery was unsuccessful and no infarct or a hemorrhagic transformation was observed. Experimental details (such as strain, sex, weight, and age, as well as the number of dead and excluded animals) are provided in the specific sections.

### 3. Permanent Middle Cerebral Artery Occlusion

All procedures were approved by and conducted according to the Danish Animal Health Care Committee (J.nr. 2019-15-0201-01620). Adult male C57BL/6 mice (21-27g, 7-8 weeks) were purchased from Taconic Ltd. (Ry, Denmark) and then group-housed in the Laboratory of Biomedicine, University of Southern Denmark under a 12 h light/dark cycle with ad libitum access to food and water. Mice were housed individually after surgery.

All surgeries were performed under Hypnorm and Dormicum anaesthesia (fentanyl citrate (0.315 mg/ml; Jansen-Cilag) and fluanisone (10 mg/ml; Jansen-Cilag, Birkerød, Denmark)), and midazolam (5 mg/ml; Hoffmann - La Roche, Hvidovre, Denmark), respectively. Focal ischemia was introduced by permanent occlusion of the distal part of the middle cerebral artery (MCA) by electrocoagulation as previously described.^71,72^ For post-operative care, 1 ml of 0.9% saline was administered s.c. and mice were transferred to a recovery room with 25°C. Mice for functional assessment were transferred to a behavioral room after 24 h. Moreover, mice were treated 3 times at eight-hour intervals with 0.001 mg/20 g buprenorphine hydrochloride (Temgesic, Reckitt & Coleman Products, UK) for postsurgical pain management, starting directly after surgery. Mice receiving sham surgery were treated in the exact same way, except that the surgery was performed without electrocoagulation of the MCA.

Mice were treated with an intraperitoneal (i.p.) injection of 175 mg/kg HOCPCA diluted in dH_2_O (10 mg/ml stock solution) or 0.9% saline for all pMCAO experiments. Two sperate studies were performed for infarct volumetric and functional analysis, where mice were allowed to survive 3 days after pMCAO, and received treatment either early (30 min, n=17/group) or at a late (3 h, n=18-19/group) timepoint after pMCAO. One study was performed for Western blot analysis: a group of mice was subjected to either sham (n=5-6/group) or pMCAO surgery (n=7-8/group) and treated 30 min post-stroke with HOCPCA or saline as stated above. These mice were allowed to survive 2 h post-pMCAO.

A total of 7 mice died after pMCAO resulting in a mortality of 6,7%, which was independent of treatment (i.e., 3 saline-treated and 2 HOCPCA-treated mice died in the early treatment group as well as one saline-treated mouse subjected to sham surgery; in the late treatment group one HOCPCA-treated mouse died). In addition, 3 mice were excluded from volumetric and functional analysis due to the lack of a visible infarct, possibly due to unsuccessful surgical occlusion or spontaneous reperfusion of the MCA.

#### 3.1. Tissue processing and infarct size assessment

For histological assessment of infarct size, mice were euthanized by cervical dislocation. Brains were extracted and immediately snap-frozen in gaseous CO_2_. Brains were stored at -80°C until further processing.

On the day of the experiment, brains were cut on a CM1860 cryostat (Leica biosystems, Germany) at -20 °C in a series of 30 μm-thick coronal sections, and cresyl violet staining was performed as described.^73^ In brief, every six sections were mounted onto Super Frost Plus slides (ThermoScientific, Slangerup, Denmark), air-dried, and passed through ascending and then descending concentrations of ethanol, followed by immersion in dH_2_0. Next, sections were stained with 0.2% cresyl violet solution for 10 min, immersed in ascending concentration of ethanol, cleared in xylene, and coverslipped with Dibutylphthalate Polystyrene Xylene (DPX) mounting media. Images were obtained and quantified with a 1.25x objective and cellSens Entry software (Olympus). Infarct volume was calculated with the following equation:

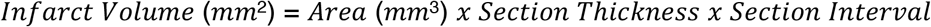

### 4. Thromboembolic stroke

Thromboemebolic stroke surgeries were performed with adult male C57BL/6 mice (10 weeks, 23-27 g), purchased from Janvier Labs (France), and subsequently group-housed in CRUB (Caen, France) under diurnal lighting conditions and free access to water and food. All experiments were performed in accordance with French ethical laws (Decree 2013-118) and European Communities Council guidelines (2010/63/EU).

Mice were anaesthetized with 5% isoflurane for induction, and anaesthesia was maintained with 2% isoflurane in a mixture of O_2_/N_2_O (30%/70%). Thromboembolic stroke surgery was performed as previously described^40^ by injection of 1μl of murine α-thrombin (1.0 IU) with a custom-made glass micropipette into the lumen of the MCA to induce *in situ* formation of a clot. Cessation of cerebral blood flow after α-thrombin injection was monitored by laser Doppler flowmetry using a fiber-optic probe (Oxford Optronix, Abington, UK). Mice received 10 mg/kg buprenorphine s.c. and local lidocaine for pain management before surgery. Mice were allowed to recover on a 37°C warm heating pad for at least 1 h post-surgery and monitored daily. Mice receiving sham surgery were treated the exact same way but were not injected with α-thrombin.

For Western blot analysis, a group of mice received either sham or stroke surgery (n=6/group), and brains were harvested 30 min post α-thrombin injection. For infarct volumetric and functional analysis after HOCPCA, tPA, HOCPCA, and t-tPA co-administration compared to vehicle, mice were allowed to survive 3 days (n=15/group). For this purpose, mice were treated with 175 mg/kg HOCPCA or 0.9% saline control 30 min post-α-thrombin injection (i.p.) as described above. In addition, a group of mice received intravenous (i.v.) injection of tPA (10 mg/kg, 10% bolus/90% infusion over 40 minutes) or the same volume of 0.9% saline starting from 20 min after α-thrombin injection via a catheter inserted into the tail vein to induce thrombolysis.

In total, 5 mice exposed to thromboembolic stroke surgery died (mortality of 8%, 3 mice died during surgery, and 2 on post-operational day 1). A total of 5 mice were excluded due to unsuccessful insertion of α-thrombin resulting in no visible infarct formation. In addition, 2 mice were excluded due to subcortical lesion formation, which results in different functional outcome. Additional animals were set up to replace these animals.

#### 4.1. Magnetic resonance imaging (MRI) analysis

Magnetic resonance imaging with a Pharmascan 7T (Bruker, Germany) was used for the determination of infarct size after thromboembolic stroke surgery 24 h post-surgery. Therefore, anesthesia was induced with 5% isoflurane and maintained with 1,5–2% isoflurane with O_2_/N_2_O (30%/70%) during the acquisitions. T2-weighted images were generated with a multi-slice multi-echo sequence (MSME, TE/TR 33 ms/2500 ms), and the volume of infarction was quantified using ImageJ (v1.53 k). Moreover, T2*-weighted images were obtained to examine if a hemorrhagic event occurred. No hemorrhage was detected.

### 5. Grip strength task

To assess functional outcome after ischemic stroke surgery, mice were tested in a grip strength task using a grip strength meter (BIO-GT3, BIOSEB, Vitrolles, France). This test takes advantage of the forepaw-grasping reflex of mice when presented with a metal bar or grid, thus quantifying muscular strength by measuring the maximum force that is required to make a mouse release its grip.

Mice were either housed in the behavior room (pMCAO) or allowed to acclimatize to the room at least 30 min prior to testing (thromboembolic stroke). Measurements were performed by the same experimenter blinded to treatment groups. The grip strength of the left and right front paw was evaluated the day before surgery (baseline), 3 days post-pMCAO as well as 1 and 3 days post-thromboembolic stroke. During the test, mice were held by the tail, dropped against a metal bar/grid, and pulled backwards horizontally until the mouse let go of the bar/grid. Each mouse was given 5 trials, and the highest force score of the 5 trials was used to calculate the grip strength asymmetry presented as a ratio of the left and right front paw. Mice were excluded if they were unable to complete the test. A total of 3 mice were excluded for pMCAO and 2 for thromboembolic stroke experiments.

### 6. Subcellular fractionation

At 2 h after pMCAO, mice were euthanized by cervical dislocation. Brains were isolated and immediately snap-frozen on crushed dry ice, and stored at -80°C. For dissection of peri-infarct cortical tissue, 1 mm-thick coronal sections were cut in the chamber of a CM1860 cryostat at -20 °C, and peri-infarct tissue was collected from the top quadrant of the left hemisphere using a tissue punch. The subcellular fractionation protocol was adapted from ^74^. In brief, tissue homogenization was achieved on ice using a tissue douncer (15 strokes) in 1 ml homogenization buffer containing 0.32 M sucrose, 1 mM NaHCO3,1mM MgCl2,10mM HEPES, pH 7.4, as well as 1,5% cOmpleteTM protease inhibitor cocktail (Roche Diagnostics), 1% phosphatase inhibitor cocktail 2 (P5726, Sigma-Aldrich) and 1% phosphatase inhibitor cocktail 3 (P0044, Sigma-Aldrich). Homogenates were centrifuged at 1000 rpm for 5 min at 4°C in a table centrifuge (5424 R, Eppendorf, Germany). The supernatant was transferred to a new tube and again centrifuged at 4°C for 5 min at 1400 rpm. After this removal of cell debris, the supernatant was centrifuged at 15 000 rpm for 15 min at 4°C, resulting in the cytosolic-enriched supernatant (S2) and membrane-enriched pellet (P2). The pellet was resuspended in 300 μl 1 mM HEPES buffer, and the supernatant was saved. Protein concentration was determined with the BCA protein assay kit (Pierce), and samples were prepared for Western blot analysis.

### 7. Western Blot

For Western blot analysis of whole-cell homogenate of peri-infarct cortical tissue after thromboembolic stroke, mice were euthanized 30 min post-stroke, and tissue was dissected as described above. Tissue was homogenized in ice-cold radioimmunoprecipitation (RIPA) buffer supplemented with 1% protease/phosphatase inhibitor cocktails (9x volume/weight) using a Bullet Blender (NextAdvance, NY, USA) and 2x 1 mm zirconium beads for 20 sec at max speed. Protein concentration was determined with the Pierce BCA protein assay, and homogenates were stored at -80°C until further analysis.

The western blot was performed and analyzed by an experimenter blinded to treatment groups. Samples were prepared on ice with a protein concentration of 0,5 μg/μl by addition of 4x Fluorescent Compatible Sample Buffer (#LC2570, Thermo Fisher) and 100 mM DTT. Samples were heated for 5 min at 95°C, sonicated, and centrifuged at 15 000 rpm for 2 min at 4°C. SDS-PAGE was performed for 40 min (200 V) with 1x Tris/glycine/SDS (25 mM Tris, 192 mM glycine, 0.1% SDS, pH 8.3) running buffer by loading 5 μg sample onto 4-20% Mini-PROTEAN® TGXTM gels (Bio-Rad) with iBrightTM Prestained Protein Ladder (#LC5615, Invitrogen). Sample order on protein gels was allocated blinded and random. Proteins were transferred to a PVDF membrane (#4561096, Biorad) using the Trans-Blot® TurboTM transfer system (Bio-Rad) (2.5A, 7min), and membranes were blocked 1x BlockerTM FL Fluorescent Blocking Buffer (#37565, Thermo Fisher) for 30 min at room temperature. The membranes were then incubated with primary antibodies followed by 3x 5 min washes in 1x tris-buffered saline (TBS) with 0.1% Tween-20 detergent (TBS-T), probed with species-specific secondary antibodies, and washed again 3x for 10 min with TBS-T. Antibodies have been diluted in 1% (w/v) BSA in TBS-T and have been reused up to four times. Image acquisition was performed with the iBright FL1500 imaging system (Invitrogen), and signals were quantified in Image Studio (Lite version 5.2).

### 8. Statistics

Statistical analysis was performed using Graphpad Prism (v.9). All data were tested for normal distributing with the Kolmogorov-Smirnov test. Data are reported as mean ± standard deviation (SD), and *in vivo* data are displayed as box and whisker graphs. Details of specific statistical tests are displayed within the figure legends. Statistical significance was accepted at p < 0.05.

## References

1. Feigin VL, Brainin M, Norrving B, et al. World Stroke Organization (WSO): Global Stroke Fact Sheet 2022. Int J Stroke. 2022;17:18–29.

2. Feigin VL, Stark BA, Johnson CO, et al. Global, regional, and national burden of stroke and its risk factors, 1990-2019: A systematic analysis for the Global Burden of Disease Study 2019. Lancet Neurol. 2021;20:1–26.

3. Grefkes C, Fink GR. Recovery from stroke: current concepts and future perspectives. Neurol Res Pract. 2020;2.

4. Fisher M, Savitz SI. Pharmacological brain cytoprotection in acute ischaemic stroke — renewed hope in the reperfusion era. Nat Rev Neurol. 2022;0123456789.

5. Hill MD, Goyal M, Menon BK, et al. Efficacy and safety of nerinetide for the treatment of acute ischaemic stroke (ESCAPE-NA1): a multicentre, double-blind, randomised controlled trial. Lancet. 2020;395:878–87.

6. Deng G, Orfila JE, Dietz RM, et al. Autonomous CaMKII Activity as a Drug Target for Histological and Functional Neuroprotection after Resuscitation from Cardiac Arrest. Cell Rep. 2017;18:1109–17.

7. Vest RS, O’Leary H, Coultrap SJ, et al. Effective post-insult neuroprotection by a novel Ca2+/ calmodulin-dependent protein kinase II (CaMKII) inhibitor. J Biol Chem. 2010;285:20675–82.

8. Leurs U, Klein AB, McSpadden ED, et al. GHB analogs confer neuroprotection through specific interaction with the CaMKIIα hub domain. Proc Natl Acad Sci. 2021;118:e2108079118.

9. Guo X, Zhou J, Starr C, et al. Preservation of vision after CaMKII-mediated protection of retinal ganglion cells. Cell. 2021;184:4299–4314.e12.

10. Rostas JAP, Spratt NJ, Dickson PW, et al. The role of Ca2+-calmodulin stimulated protein kinase II in ischaemic stroke - A potential target for neuroprotective therapies. Neurochem Int. 2017;107:33–42.

11. Rosenberg OS, Deindl S, Comolli LR, et al. Oligomerization states of the association domain and the holoenyzme of Ca2+ / CaM kinase II. FEBS J. 2006;273:682–94.

12. Paul De Koninck, Schulman H. Sensitivity of CaM kinase II to the frequency of Ca2+ oscillations. Science. 1998;279:227–30.

13. Cook SG, Buonarati OR, Coultrap SJ, et al. CaMKII holoenzyme mechanisms that govern the LTP versus LTD decision. Sci Adv. 2021;7:eabe2300.14.

14. Bayer KU, De Koninck P, Leonard AS, et al. Interaction with the NMDA receptor locks CaMKII in an active conformation. Nature. 2001;411:801–5.

15. Barria A, Muller D, Derkach V, et al. Regulatory phosphorylation of AMPA-type glutamate receptors by CaM-KII during long-term potentiation. Science. 1997;276:2042–5.

16. Bayer KU, Schulman H. CaM Kinase: Still Inspiring at 40. Neuron. 2019;2.

17. Hell JW. CaMKII: Claiming center stage in postsynaptic function and organization. Neuron. 2014;81:249–65.

18. Aronowski J, Grotta JC. Ca2+/ calmodulin-dependent protein kinase II in postsynaptic densities after reversible cerebral ischemia in rats. Brain Res. 1996;709:103–10.

19. Matsumoto S, Shamloo M, Matsumoto E, et al. Protein Kinase C-γ and Calcium/Calmodulin-Dependent Protein Kinase II-α Are Persistently Translocated to Cell Membranes of the Rat Brain during and after Middle Cerebral Artery Occlusion. J Cereb Blood Flow Metab. 2004;24:54–61.

20. Ameen SS, Griem-Krey N, Dufour A, et al. N-Terminomic Changes of Neurons During Excitotoxicity Reveals Pathological Proteolytic Events Associated with Synaptic Dysfunctions and Potential Targets for Neuroprotection. bioRxiv. 2020;

21. Coultrap SJ, Vest RS, Ashpole NM, et al. CaMKII in cerebral ischemia. Acta Pharmacol Sin. 2011;32:861–72.

22. Waxham MN, Grotta JC, Silva AJ, et al. Ischemia-induced neuronal damage: a role for calcium/calmodulin-dependent protein kinase II. J Cereb Blood Flow Metab. 1996;16:1–6.

23. Wellendorph P, Høg S, Greenwood JR, et al. Novel cyclic γ-hydroxybutyrate (GHB) analogs with high affinity and stereoselectivity of binding to GHB sites in rat brain. J Pharmacol Exp Ther. 2005;315:346–51.

24. Ottani A, Saltini S, Bartiromo M, et al. Effect of γ-hydroxybutyrate in two rat models of focal cerebral damage. Brain Res. 2003;986:181–90.

25. Sadasivan S, Maher TJ, Quang LS. γ-hydroxybutyrate (GHB), γ-butyrolactone (GBL), and 1,4-butanediol (1,4-BD) reduce the volume of cerebral infarction in rodent transient middle cerebral artery occlusion. Ann N Y Acad Sci. 2006;1074:537–44.

26. Gao B, Kilic E, Baumann CR, et al. γ-hydroxybutyrate accelerates functional recovery after focal cerebral ischemia. Cerebrovasc Dis. 2008;26:413–9.

27. Kaupmann K, John F, Wellendorph P, et al. Specific γ-hydroxybutyrate-binding sites but loss of pharmacological effects of γ-hydroxybutyrate in GABAB(1)-deficient mice. Neuroscience. 2003;18:2722–30.

28. Thiesen L, Kehler J, Clausen RP, et al. In vitro and in vivo evidence for active brain uptake of the GHB analog HOCPCA by the monocarboxylate transporter subtype 1. J Pharmacol Exp Ther. 2015;354:166–74.

29. Vogensen SB, Marek A, Bay T, et al. New synthesis and tritium labeling of a selective ligand for studying high-affinity γ-hydroxybutyrate (GHB) binding sites. J Med Chem. 2013;56:8201–8205.

30. Pellicena P, Schulman H. CaMKII inhibitors: From research tools to therapeutic agents. Front Pharmacol. 2014;5:1–10.

31. STAIR. Recommendations for standards regarding preclinical neuroprotective and restorative drug development. Stroke. 1999;30:2752–8.

32. Fisher M, Feuerstein G, Howells DW, et al. Update of the stroke therapy academic industry roundtable preclinical recommendations. Stroke. 2009;40:2244–50.

33. McCabe C, Arroja MM, Reid E, et al. Animal models of ischaemic stroke and characterisation of the ischaemic penumbra. Neuropharmacology. 2017;1–9.

34. Sommer CJ. Ischemic stroke: experimental models and reality. Acta Neuropathol. 2017;133:245–61.

35. Hossmann KA. The two pathophysiologies of focal brain ischemia: Implications for translational stroke research. J Cereb Blood Flow Metab. 2012;32:1310–6.

36. Bach A, Clausen BH, Moller M, et al. A high-affinity, dimeric inhibitor of PSD-95 bivalently interacts with PDZ1-2 and protects against ischemic brain damage. Proc Natl Acad Sci. 2012;109:3317–22.

37. Matsumoto S, Shamloo M, Isshiki A, et al. Persistent Phosphorylation of Synaptic Proteins Following Middle Cerebral Artery Occlusion. J Cereb Blood Flow Metab. 2002;22:1107–13.

38. Buonarati OR, Cook SG, Goodell DJ, et al. CaMKII versus DAPK1 Binding to GluN2B in Ischemic Neuronal Cell Death after Resuscitation from Cardiac Arrest. Cell Rep. 2020;30:1–8.e4.

39. Miller SG, Kennedy MB. Distinct forebrain and cerebellar isozymes of type II Ca2+/calmodulin-dependent protein kinase associate differently with the postsynaptic density fraction. J Biol Chem. 1985;260:9039–46.

40. Orset C, Macrez R, Young AR, et al. Mouse model of in situ thromboembolic stroke and reperfusion. Stroke. 2007;38:2771–8.

41. Gauberti M, Martinez de Lizarrondo S, Orset C, et al. Lack of secondary microthrombosis after thrombin-induced stroke in mice and non-human primates. J Thromb Haemost. 2014;12:409–14.

42. Molina CA, Montaner J, Abilleira S, et al. Timing of spontaneous recanalization and risk of hemorrhagic transformation in acute cardioembolic stroke. Stroke. 2001;32:1079–84.

43. De Lizarrondo SM, Gakuba C, Herbig BA, et al. Potent thrombolytic effect of N-acetylcysteine on arterial thrombi. Circulation. 2017;136:646–60.

44. Drieu A, Buendia I, Levard D, et al. Immune Responses and Anti-inflammatory Strategies in a Clinically Relevant Model of Thromboembolic Ischemic Stroke with Reperfusion. Transl Stroke Res. 2019;

45. Rostas JAP, Hoffman A, Murtha LA, et al. Ischaemia- and excitotoxicity-induced CaMKII-mediated neuronal cell death: The relative roles of CaMKII autophosphorylation at T286 and T253. Neurochem Int. 2017;104:6–10.

46. Cruzalegui FH, Kapilofft MS, Morfint J, et al. Regulation of intrasteric inhibition of the multifunctional calcium/calmodulin-dependent protein kinase. PNAS. 1992;89:12127–31.

47. Pi HJ, Otmakhov N, Lemelin D, et al. Autonomous CaMKII can promote either long-term potentiation or long-term depression, depending on the state of T305/T306 phosphorylation. J Neurosci. 2010;30:8704–9.

48. Hayashi Y, Shi S, Esteban JA, et al. Driving AMPA Receptors into Synapses by LTP and CaMKII : Requirement for GluR1 and PDZ Domain Interaction. Science. 2000;287:2262–8.

49. Kwiatkowski AP, King MM. Autophosphorylation of the Type II Calmodulin-Dependent Protein Kinase Is Essential for Formation of a Proteolytic Fragment with Catalytic Activity. Implications for Long-Term Synaptic Potentiation. Biochemistry. 1989;28:5380–5.

50. Hossain MI, Roulston CL, Kamaruddin MA, et al. A truncated fragment of Src protein kinase generated by calpain-mediated cleavage is a mediator of neuronal death in excitotoxicity. J Biol Chem. 2013;288:9696–709.

51. Wong MH, Samal AB, Lee M, et al. The KN-93 Molecule Inhibits Calcium/Calmodulin-Dependent Protein Kinase II (CaMKII) Activity by Binding to Ca2+/CaM. J Mol Biol. 2019;431:1440–59.

52. Vest RS, Davies KD, O’Leary H, et al. Dual mechanism of a natural CaMKII inhibitor. Mol Biol Cell. 2007;18:5024–33.

53. Nassal D, Gratz D, Hund TJ. Challenges and opportunities for therapeutic targeting of calmodulin kinase II in heart. Front Pharmacol. 2020;11:1–13.

54. Ashpole NM, Chawla AR, Martin MP, et al. Loss of calcium/calmodulin-dependent protein kinase II activity in cortical astrocytes decreases glutamate uptake and induces neurotoxic release of ATP. J Biol Chem. 2013;288:14599–611.

55. Ye J, Das S, Roy A, et al. Ischemic Injury-Induced CaMKIIδ and CaMKIIγ Confer Neuroprotection Through the NF-κB Signaling Pathway. Mol Neurobiol. 2019;56:2123–36.

56. Buard I, Coultrap SJ, Freund RK, et al. CaMKII “Autonomy” Is Required for Initiating But Not for Maintaining Neuronal Long-Term Information Storage. J Neurosci. 2010;30:8214–20.

57. Sloutsky R, Dziedzic N, Dunn MJ, et al. Heterogeneity in human hippocampal CaMKII transcripts reveals allosteric hub-dependent regulation. Sci Signal. 2020;13:1–13.

58. Bhattacharyya M, Stratton MM, Going CC, et al. Molecular mechanism of activation-triggered subunit exchange in Ca2+/calmodulin-dependent protein kinase II. Elife. 2016;5:1–32.

59. Stratton M, Lee IH, Bhattacharyya M, et al. Activation-triggered subunit exchange between CaMKII holoenzymes facilitates the spread of kinase activity. Elife. 2014;3:e01610.

60. Karandur D, Bhattacharyya Rm, Xia Z, et al. Breakage of the oligomeric CaMKII hub by the regulatory segment of the kinase. Elife. 2020;9:1–31.

61. Bhattacharyya M, Lee YK, Muratcioglu S, et al. Flexible linkers in CaMKII control the balance between activating and inhibitory autophosphorylation. Elife. 2020;9.

62. Strack S, Barban MA, Wadzinski BE, et al. Differential inactivation of postsynaptic density-associated and soluble Ca2+/calmodulin-dependent protein kinase II by protein phosphatases 1 and 2A. J Neurochem. 1997;68:2119–28.

63. Chalmers NE, Yonchek J, Steklac KE, et al. Calcium/Calmodulin-Dependent Kinase (CaMKII) Inhibition Protects Against Purkinje Cell Damage Following CA/CPR in Mice. Mol Neurobiol. 2020;57:150–8.

64. Harada K, Maekawa T, Tsuruta R, et al. Hypothermia inhibits translocation of CaM kinase II and PKC-α, β, γ isoforms and fodrin proteolysis in rat brain synaptosome during ischemia-reperfusion. J Neurosci Res. 2002;67:664–9.

65. Gurd JW, Rawof S, Zhen Huo J, et al. Ischemia and status epilepitcus result in enhanced phosphorylation of calcium and calmodulin-stimulated protein kinase II on threonine 253. Brain Res. 2008;1218:158–65.

66. Hu B-R, Kurihara J, Wieloch T. Persistent Translocation and Inhibition of Ca2+/Calmodulin-Dependent Protein Kinase II in the Crude Synaptosomal Fraction of the Vulnerable Hippocampus Following Hypoglycemia. J Neurochem. 1995;64:1361–9.

67. Mengesdorf T, Althausen S, Mies G, et al. Phosphorylation state, solubility and activity of Calcium/Calmodulin-Dependent Protein Kinase IIa in Transient Focal Ischemia in Mouse Brain. Neurochem Res. 2002;27:477–84.

68. Skelding KA, Spratt NJ, Fluechter L, et al. αCaMKII is differentially regulated in brain regions that exhibit differing sensitivities to ischemia and excitotoxicity. J Cereb Blood Flow Metab. 2012;32:2181–92.

69. McCabe C, Arroja MM, Reid E, et al. Animal models of ischaemic stroke and characterisation of the ischaemic penumbra. Neuropharmacology. 2018;134:169–77.

70. Durand A, Chauveau F, Cho TH, et al. Spontaneous Reperfusion after In Situ Thromboembolic Stroke in Mice. PLoS One. 2012;7.

71. Lambertsen KL, Gregersen R, Drøjdahl N, et al. A specific and sensitive method for visualization of tumor necrosis factor in the murine central nervous system. Brain Res Protoc. 2001;7:175–91.

72. Lambertsen KL, Clausen BH, Babcock AA, et al. Microglia protect neurons against ischemia by synthesis of tumor necrosis factor. J Neurosci. 2009;29:1319–30.

73. Griem-Krey N, Klein AB, Herth MM, et al. Autoradiography as a simple and powerful method for visualization and characterization of pharmacological targets. J Vis Exp. 2019;e58879.

74. Kool MJ, Onori MP, Borgesius NZ, et al. CAMK2-dependent signaling in neurons is essential for survival. J Neurosci. 2019;1341–18.

