## Supporting information for "The GHB analogue HOCPCA improves sensorimotor function after MCAO via CaMKIIα"

### Supplementary figures

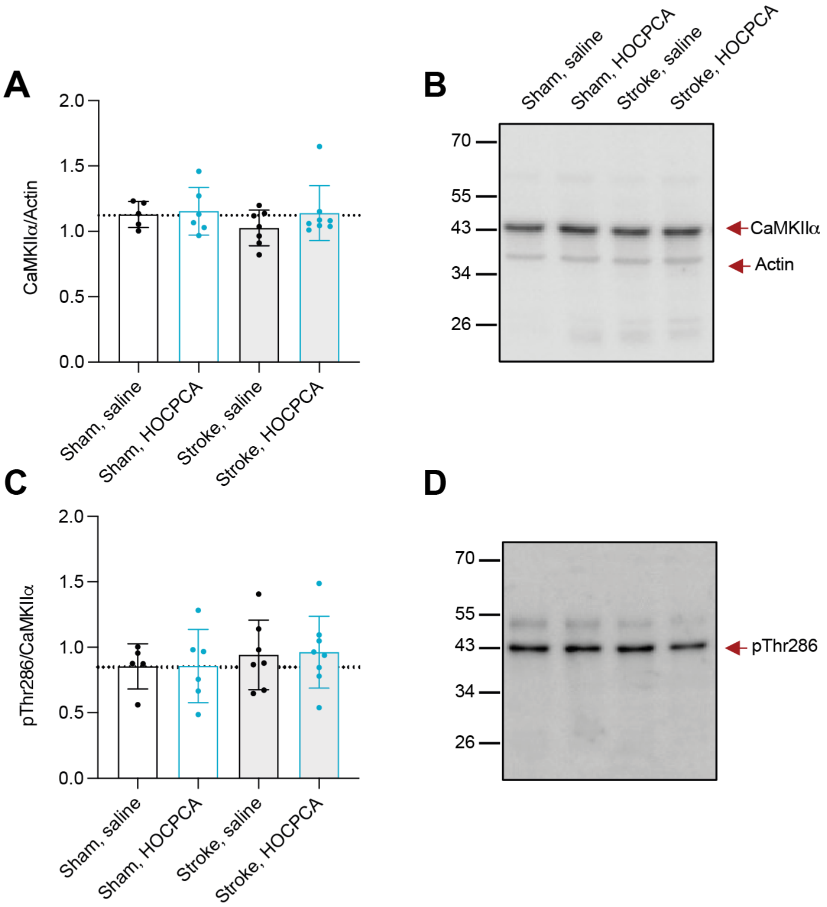


**Supplementary figure 1: Effect of HOCPCA on CaMKIIα expression and Thr286 autophosphorylation after pMCAO.** Adult male mice were subjected to sham (*n* = 5-6/group) or pMCAO-surgery (*n* = 7-8/group), treated with 175 mg/kg HOCPCA or saline control and brains were harvested 2 h post-stroke. Whole-cell homogenates from peri-infarct tissue were probed for (*A*) total CaMKIIα or (*C*) pThr286 and normalized to Actin or total CaMKIIα expression. (*B,D*) Representative Western blots.

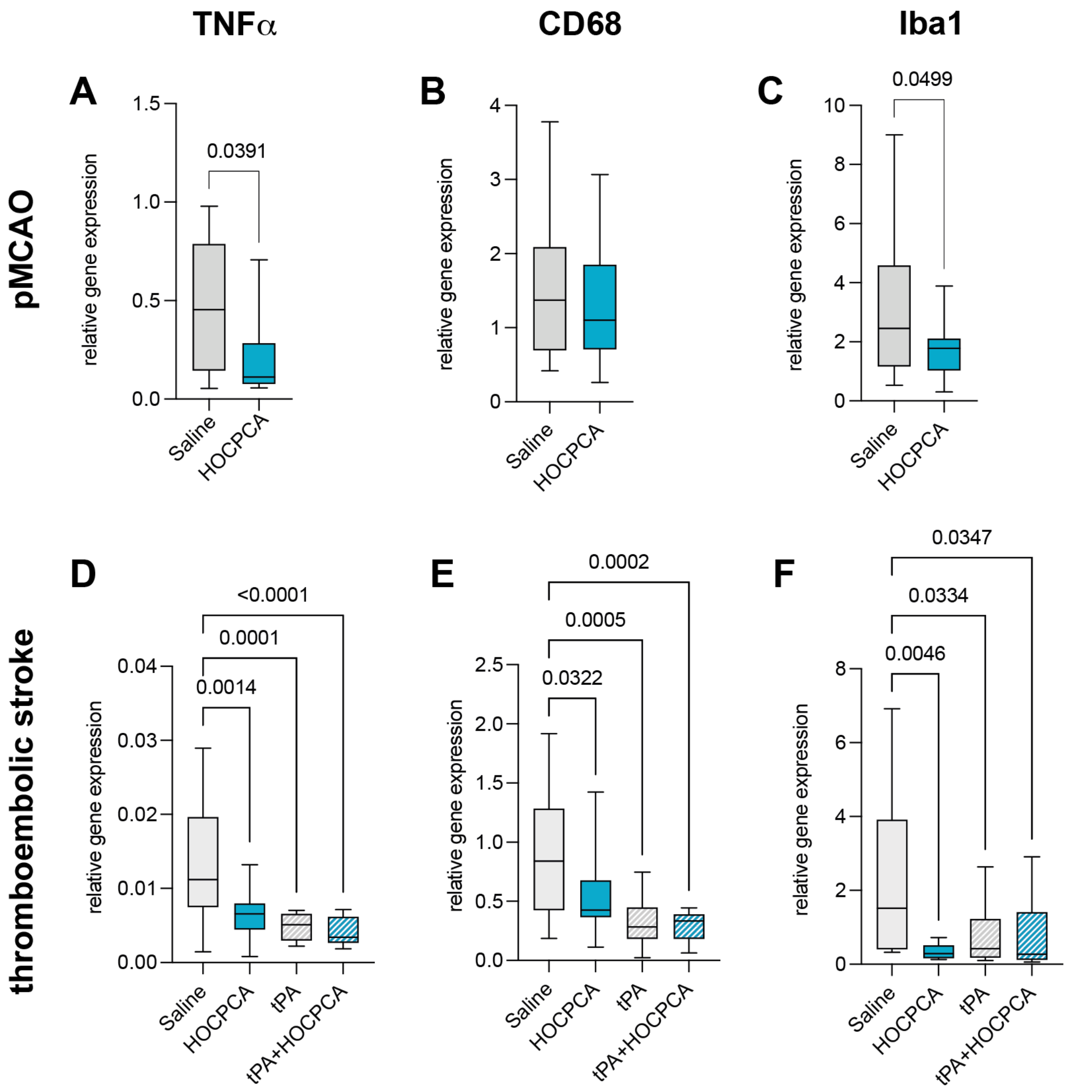


**Supplementary figure 2: Treatment effect on mRNA expression of inflammatory markers after pMCAO and thromboembolic stroke.** (*A-F)* Ipsilateral cortex was collected 3 days-stroke for qPCR analysis and is presented relative to the mRNA expression of the reference gene *Sdha*. mRNA Expression of (*A,D*) *Tnfa* (TNFα) (*B,F*) *Cd68* (CD68) and (*C,F*) *Aif1* (Iba1). (*A-C*) Adult male mice receiving pMCAO surgery and treatment at 30 min post-stroke with 175 mg/kg HOCPCA or saline (*n* = 17/group; two-tailed Student's *t*-test). (*D*-F) Adult male mice receiving thromboembolic stroke surgery were treated either with saline, tPA (intravenous, 10 mg/kg, 10% bolus/90% infusion over 40 minutes starting from 20 min after occlusion), HOCPCA (intraperitoneal, at 30 min post-stroke) or a combination of tPA and HOCPCA (*n* = 15/group; one-way ANOVA, post hoc Tukey's test) (Box plot (boxes, 25–75%; whiskers, minimum and maximum; lines, median).

### Supplementary materials and methods

*1. Reverse transcription quantitative polymerase chain reaction (RT-qPCR)*

To determine mRNA expression changes of inflammatory markers, ipsilateral cortical tissue from the ischemic area was collected 3 days post-stroke. For pMCAO experiments, tissue was collected while processing for infarct volumetric analysis (see section 3.1). For thromboembolic stroke experiments, mice were euthanized by cervical dislocation, brains were dissected out rapidly, snap-frozen on crushed dry-ice and stored at -80 °C until further processing. On the day of the assay, brains were dissected using a CM1860 cryostat (Leica) at - 20 °C and ischemic cortical tissue was collected using a tissue punch.

Tissue was homogenized using RLT plus Lysis buffer (Qiagen) with zirconium oxide beads (2 x 2 mm) in a Bullet Blender, and RNA extraction was performed using the RNeasy Plus Mini kit (#74136, Qiagen) according to the manufacturer's instructions with genomic DNA (gDNA) eliminator columns. Total RNA amounts were quantified along with quality evaluation by spectrophotometrical determination of 260/280 and 230/260 values with a NanoDrop 2000 (Thermo Fisher). Complementary DNA (cDNA) was synthesized by reverse transcription with 500 ng of RNA using qScript® cDNA Supermix (Quanta Bio) on an Eppendorf Mastercycler Personal PCR machine (25 °C for 5 min, 42 °C for 30 min, 85 °C for 5 min). The cDNA was diluted with nuclease free water to approximately 100 ng/µl and stored at -20 °C until RT-qPCR was performed. RT-qPCR samples were prepared by combining cDNA, Power SYBR green Master Mix (2x) (Applied Biosystems), forward and reverse primers (**Suppl. Table 1**), and performed by 10 min heating to 95 °C followed by 40 PCR cycles of 15 s at 95 °C and 60 s at 60 °C on a Stratagene Mx3005P (Agilent Technologies), according to the manufactures protocol of the Master Mix. Dissociation curve analysis was carried out at 60 s at 95 °C, 30 s at 55 °C and 30 s at 90 °C to verify the presence of unique amplification products, and primers were produced by TAG Copenhagen. If possible, primer sequences were designed to span exon-exon junctions. MxPro qPCR Software (Agilent Technologies) was used to determine the cycle threshold (Ct), and the relative gene expression compared to a reference gene (*Sdha*) were calculated with the following equation, as described previously:[1]

$$\Delta C_t = 2^{(C_{t,reference} - C_{t,target})}$$

**Supplementary table 1: Primer sequences for RT-qPCR.**

| Gene | Primer Sequence |
|---|---|
| *Sdha* | F: 5' GGAACACTCCAAAAACAGACCT 3'<br>R: 5' CCACCACTGGGTATTGAGTAGAA 3' |
| *Tnfa* | F: 5' GCCTCCCTCTCATCAGTTCTAT 3'<br>R: 5' TTTGCTACGACGTGGGCTA 3' |
| *Cd68* | F: 5' GGGGCTCTTGGGAACTACAC 3'<br>R: 5' GTACCGTCACAACCTCCCTG 3' |
| *Aif1* | F: 5' GTCCTTGAAGCGAATGCTGG 3'<br>R: 5' CATTCTCAAGATGGCAGATC 3' |

### Supplementary reference


1. Schmittgen TD, Livak KJ. Analyzing real-time PCR data by the comparative CT method. *Nat Protoc*. 2008;3:1101–8.
